# Hemisynthetic derivatives of the natural alkaloid trilobine are fast-acting antimalarial compounds with sustained activity in multi-drug resistant *P. falciparum* isolates

**DOI:** 10.1101/2022.08.30.505923

**Authors:** Flore Nardella, Irina Dobrescu, Haitham Hassan, Fabien Rodrigues, Sabine Thiberge, Liliana Mancio Silva, Ambre Tafit, Corinne Jallet, Véronique Cadet-Daniel, Stéphane Goussin, Audrey Lorthiois, Yoann Menon, Nicolas Molinier, Dany Pechalrieu, Christophe Long, François Sautel, Mariette Matondo, Magalie Duchateau, Guillaume Medard, Benoit Witkowski, Artur Scherf, Ludovic Halby, Paola B. Arimondo

## Abstract

Malaria eradication requires the development of new drugs to combat drug-resistant parasites. The search for new chemical scaffolds that target novel pathways of the human malaria parasite *Plasmodium falciparum* is of highest priority. We identified bisbenzylisoquinoline alkaloids isolated from *Cocculus hirsutus*. (trilobine derivatives) as active in the nanomolar range against *P. falciparum* blood stages. Synthesis of a library of 94 hemi-synthetic derivatives allowed us to identify compound **84** that kills multi-drug resistant clinical isolates in the nanomolar range (median IC_50_ ranging from 35-88nM). Efforts were made to obtain compounds with significantly improved preclinical properties. Out of those, compound **125** delays the onset of parasitemia in *P. berghei* infected mice and inhibits *P. falciparum* transmission stages *in vitro* (culture assays) and *in vivo* using membrane feeding assay in the *Anopheles stephensi* vector. Compound **125** also impairs *P. falciparum* development in sporozoite-infected hepatocytes, in the low micromolar range. Finally, we used a chemical pull-down strategy to identify potential protein targets of this chemical family. Mass spectrometry analysis identified the parasite interactome with trilobine derivatives, identifying protein partners belonging to metabolic pathways that have not been previously targeted by antimalarial drugs or implicated in drug-resistance mechanisms.

## Introduction

Malaria, a vector borne disease, continues to cause morbidity and mortality with more than 627,000 annual deaths according to the World Health Organisation (WHO)^1^. This disease is caused by an apicomplexan protozoan parasite belonging to the genus *Plasmodium*. Among the five *Plasmodium spp* that infect humans, *Plasmodium falciparum* is by far responsible for most severe disease and deaths. Treatment relies on artemisinin derivatives used alone (severe malaria) or in combination (uncomplicated malaria) with chloroquine derivatives (amodiaquine, piperaquine, pyronaridine), amino-alcohols (mefloquine, lumefantrine) or antifolates (sulfadoxine, pyrimetamine). These treatments are highly efficient but resistances to drugs of these combinations, that arose in South-East Asia, complicate disease management and threatens elimination^2^. Most commonly used antimalarial medications are derived or inspired by natural products, like quinine, artemisinin, atovaquone, or doxycycline^3^. Natural products have been an incredible source of inspiration in drug discovery and fills a chemical space that has never been reached by chemical synthesis^4^.

Here, we screened an in-house chemical library of hemisynthetic derivatives of a natural product called trilobine (bisbenzylisoquinoline alkaloid) obtained from *Cocculus hirsutus*. Activity of the library was tested against *P. falciparum* asexual blood stage parasites, the stage responsible for all clinical symptoms. Hits belonging to the natural product family showed killing activity in the nanomolar range against *P. falciparum*. Structural modifications of the lead compound improved the activity against the disease-causing stage of malaria, but also in the stages that transmit the disease. This compound has a sustained activity in multi-drug resistant clinical isolates originating from Cambodia and is fast-acting. Efforts were made to develop derivatives with an optimized *in vivo* efficacy in an experimental mouse malaria model and to decipher the protein target of these natural product derivatives.

## Results

### Natural product trilobine hemisynthetic derivatives are active against *P. falciparum* asexual blood stage

Screening of an in-house chemical library that targets distinct human epigenetic factors^5,6^ identified hits against asexual blood stages of *P. falciparum*, belonging to the double bridge bisbenzylisoquinoline family of alkaloids found in *Cocculus hirsutus*. We measured the activity of 94 natural and hemisynthetic derivatives and found compound **84** as being the most active (**Figure 1A**), with a mean IC_50_ of 130nM. This compound was then chosen as the lead compound. Hemi-synthesis of **84** from the trilobine natural analogue cocsuline (**4**) is described in **Figure 1B**.

**Figure 1.**
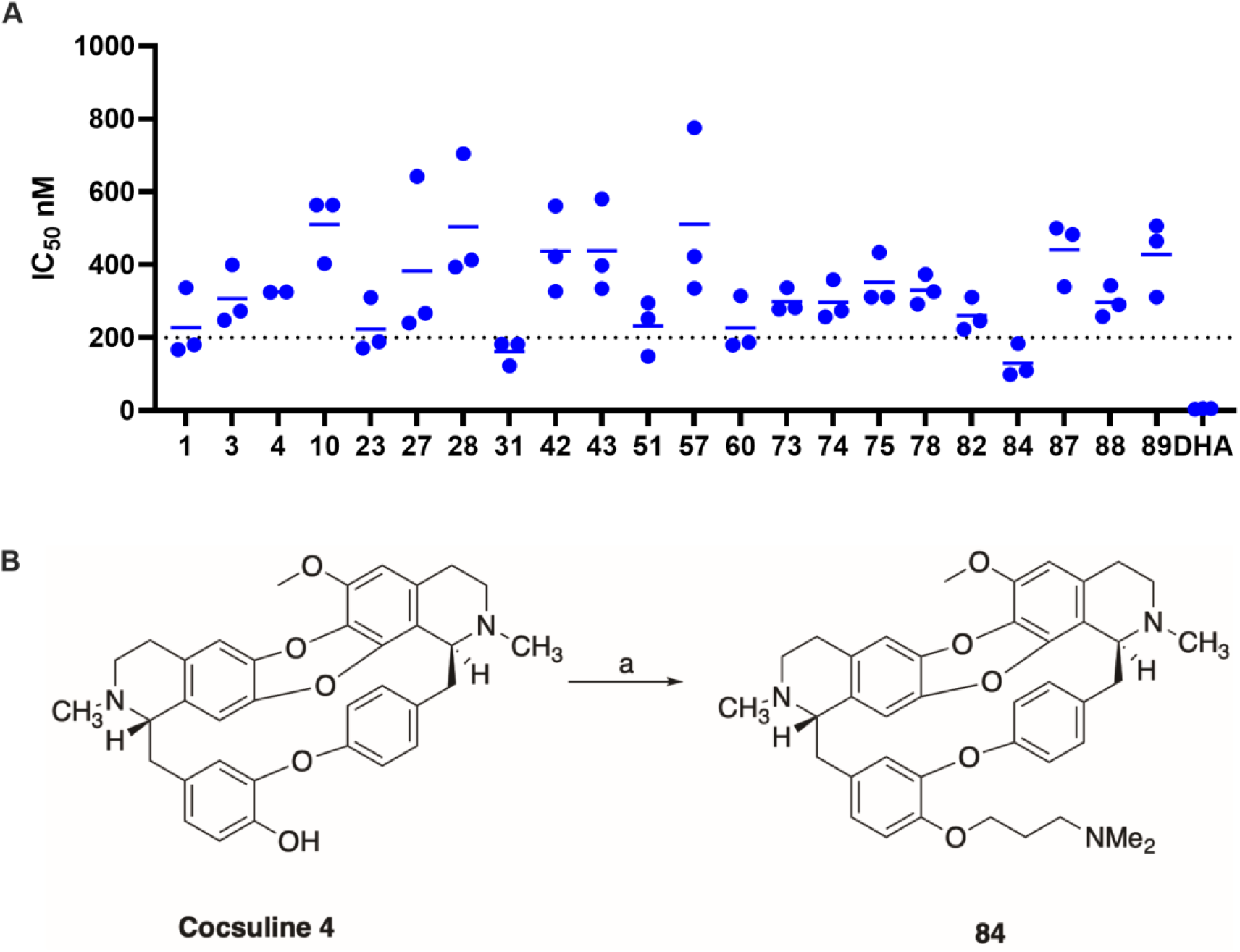
IC_50_ of the most potent compounds out of the 94 derivatives of the chemical series and hemi-synthesis of compound 84. **A**. IC_50_ was measured in triplicate using *P. falciparum* NF54 strain, in three independent experiments. Growth was determined using the SYBR-Green I assay after 72h incubation with the compounds. Points represent individual IC_50_ value (in nM) measured in each experiment and lines represent the mean. Compound **84** is the most active with a mean IC_50_ of 130 nM and was selected as the lead compound. **B**. Hemi-synthesis of **84** starting from natural compound cocsuline (**4**): *i*) NaH, DMF, 0°C then RT, 30 min; *ii*) 3-dimethylaminopropylchloride, KI, 80°C, 15h.

### 84 is a fast-acting compound and is active throughout the asexual blood cycle

Compound **84** was tested for its stage-specific activity in the asexual blood cycle (**Figure 2A**). Tightly synchronized *P. falciparum* (NF54) parasites were exposed to concentrations corresponding to 3x or 10x the IC_50_ of **84**. Parasites were treated for 6h windows covering the entire 48h cycle. Survival was assessed by counting parasites in a thin blood smear 72h post-invasion (hpi). Compound **84** is active throughout the asexual cell cycle, with the highest killing activity between 18 and 42hpi. Importantly, **84** is active in the early ring stage (0-6hpi), with less than 5% surviving parasites at 10x the IC_50_ and about 20% survival at 3x the IC_50_. As 6h of incubation were sufficient to obtain efficient parasite killing, we assessed the speed of action of **84** using the Parasite Reduction Rate assay (PRR) developed by Sanz et al^7^. We found that **84** is as fast acting as dihydroartemisinin (DHA), the active metabolite of artemisinin derivatives (**Figure 2B**).

**Figure 2.**
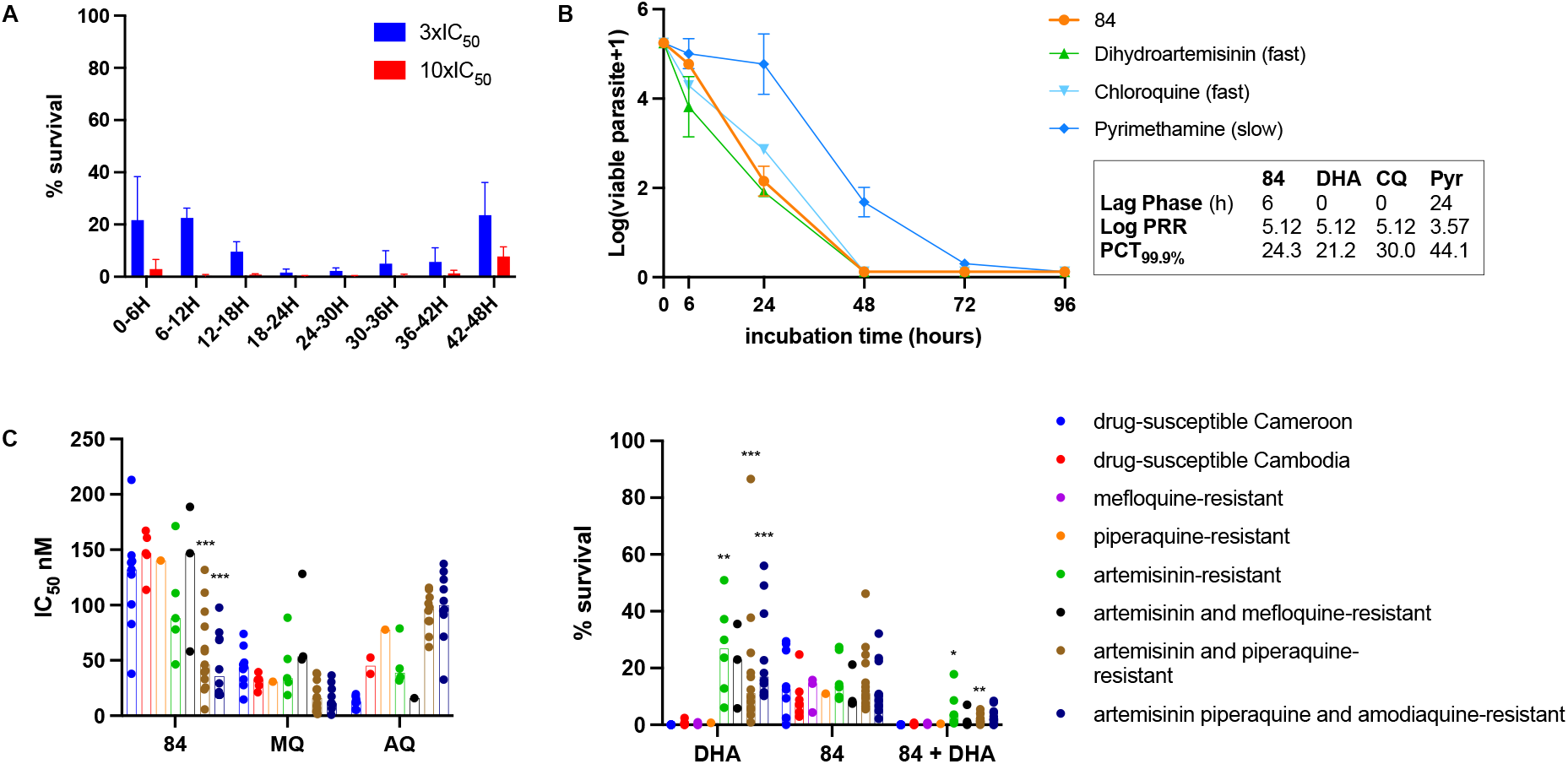
Activity of compound 84 in *P. falciparum* asexual blood stages. **A**. Stage-specific activity in *P. falciparum* NF54 asexual cell cycle. Tightly synchronized parasites (0-3h post-invasion) were incubated with **84** for 6h, for 8 consecutive 6h periods covering the 48h cell cycle, at two different concentrations (3x and 10x the IC_50_). Survival rate was evaluated 72h after synchronization (n=3 independent experiments). **B**. *In vitro* parasite reduction rate (PRR). *P. falciparum* culture containing mostly ring and mono-infected red blood cells (RBC) were incubated with **84** or reference drugs (dihydroartemisinin, chloroquine and pyrimethamine) for different time windows (6h, 24h, 48h, 72h and 96h), washed, serially diluted in fresh RBC and allowed to grow for 3 weeks, then growth was assessed using the SYBR-Green I assay^7^. N=1 experiment run in duplicate. **C**. Activity against *P. falciparum* multi-drug resistant field isolates adapted to culture. **Left graph:** IC_50_s (dots) of **84** and two reference drugs (mefloquine, MQ and desethlyamodiaquine, AQ) obtained against Cameroonian and Cambodian field isolates are plotted versus the type of resistance carried by the isolate. **Right graph:** Percentage of survival using the Ring-stage Survival Assay (RSA). Tightly synchronized ring-stage parasites (0-3h post-invasion) were incubated with 700nM dihydroartemisinin (DHA), 3x the IC_50_ of **84** (equivalent to 400nM), or the combination of both, for 6h. After 6 hours of incubation, the drugs are washed out and the parasites are allowed to grow until 72h post-synchronization. Parasite survival is assessed by counting parasitemia in 10,000 RBC in a thin blood smear, in comparison to a DMSO treated control^8^. Statistics = multiple Mann-Whitney tests in comparison with the *drug-sensitive Cambodia* condition (if discovery: ***p value<0.001; ** p value<0.01; *p value<0.05).

### Compound 84 has a sustained activity in *P. falciparum* multi-drug resistant field isolates adapted to culture

Since compound **84** is active in early ring stages and is fast-acting, we assessed its activity against multi-drug resistant field isolates, notably isolates resistant to artemisinin (**Figure 2C**). Using IC_50_ as a proxy, **84** has a similar IC_50_ in the laboratory strain NF54 and drug-sensitive isolates originating from Cameroon and Cambodia (median of 132.1 and 146.9nM, respectively). Interestingly, isolates resistant to artemisinin have an increased sensitivity to **84** (median IC_50_ of 88.2nM) especially when they carry additional resistance to piperaquine and amodiaquine, with median IC_50_ values dropping to 46.3 and 35.7nM respectively (**Figure 2C**, left panel). In the Ring-stage Survival Assay^8^, dihydroartesiminin (DHA) shows, as expected, a difference in survival between artemisinin-susceptible and artemisinin-resistant isolates (DHA concentration 200 times the IC_50_). Compound **84** shows no difference in survival, whichever the resistance background of the isolates, when used at a concentration of 3 times the IC_50._ The observed survival values correspond to the value obtained in the laboratory line NF54 (**Figure 2A**). Notably, **84** combined with DHA reduces almost to zero the survival of artemisinin resistant strains (**Figure 2C**, right), indicating a synergistic effect.

### Preclinical studies reveal that 84 binds to human plasma proteins

To assess compound **84** *in vivo*, preclinical studies were realized to test the cytotoxicity, microsomal stability and plasma binding properties. Microsomal stability half-life of **84** was higher than 60min and the cytotoxicity in HepG2 and MV411 cell-lines resulted in a selectivity index superior to 75. These results represent suitable preclinical values for drug development. However, **84** was found highly bound to plasma protein (>99% in both human and mice). This means that compound **84** would sequester in the blood to host plasma proteins, and that only a very minor fraction would be able to reach the target in infected host cell. To overcome this problem, efforts were made to decrease the plasma binding by synthetizing modified derivatives of **84** (**Figure 3A**).

**Figure 3.**
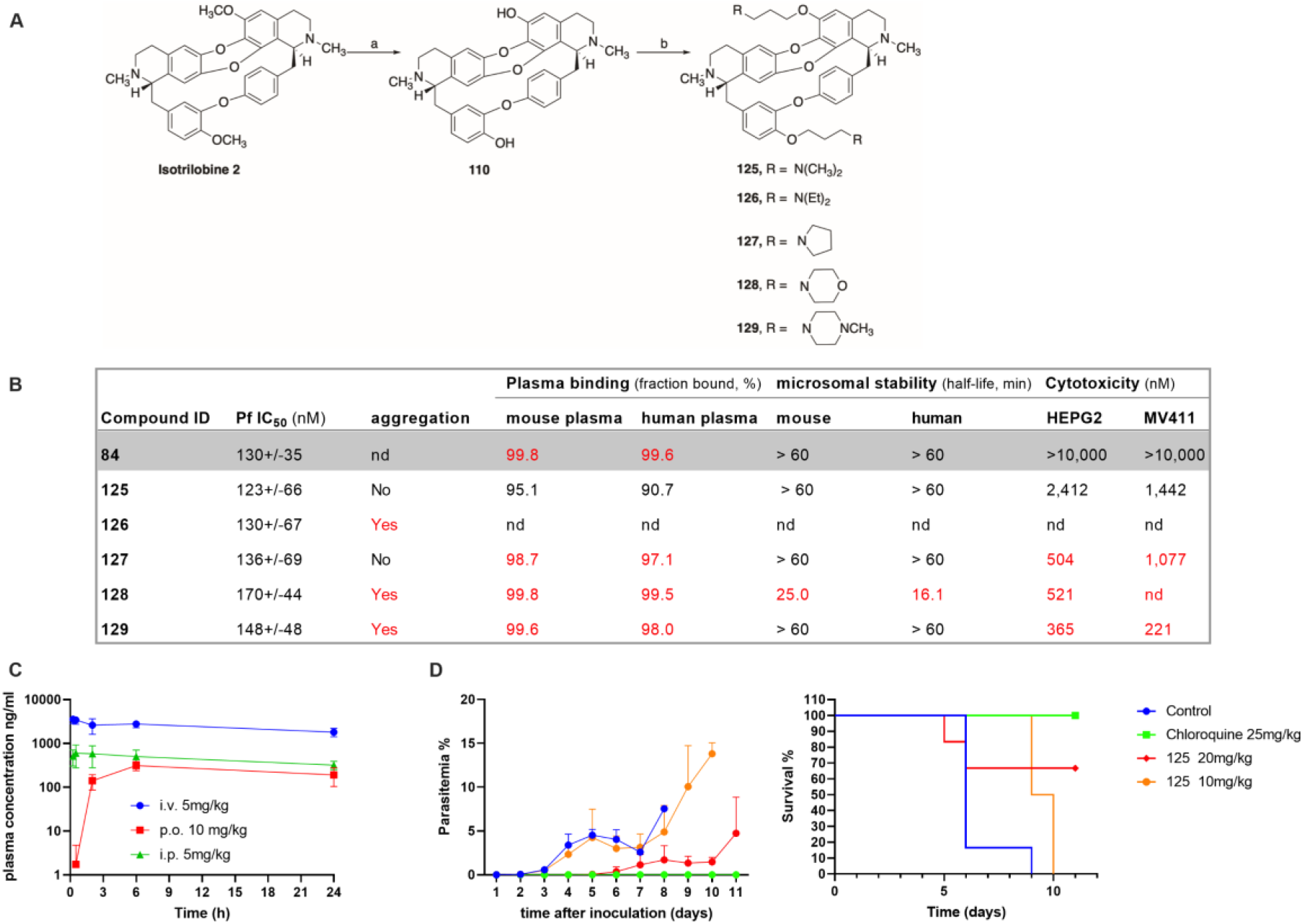
Compound 84 and its derivatives: towards the best candidate for *in vivo* studies. **A**. General synthesis of compounds **125-129. a)** *i*) NaH, EtSH, DMF, 0°C, 30 min; *ii*) isotrilobine **2**, 150°C, 30 min; b) *i*) NaH, DMF, 0°C, 30 min; *ii*) Desired chloro-derivative, 80°C, 15h **a)** ETSH, DMF, 150°C, 30min, 89%; **b)** RCl, NaH, DMF, Ar, 80°C, ON, 25-45%. **B**. Table reporting the mean IC_50_ measured against *P. falciparum*, the solubility of the compound, the plasma binding fraction in mouse and human plasma, the microsomal stability measured in mouse and human microsomes and the cytotoxicity measured in two human cell lines (HepG2 and MV411), for **84** and its derivatives. **C**. Pharmacokinetic profile of compound **125** after a single dose of compound, administered either intravenously (*i*.*v*., 5mg/kg), *per os* (*p*.*o*., 10mg/kg) or intraperitoneally (*i*.*p*., 5mg/kg). Plasmatic concentration is represented in ng/mL versus time (5 timepoints over 24h). **D**. *In vivo* activity of **125** in C57BL/6 mice infected with *P. berghei* ANKA. Mice were treated *i*.*p*., 2h after parasite *i*.*v*. inoculation, for 4 days, with a daily regimen of 20mg/kg or 10mg/kg of **125** or 25mg/kg of chloroquine or the vehicle control. Parasitemia and survival were followed over 11 days.

### Chemical synthesis of novel derivatives and *in vitro* activity

Selective double demethylation of the two methoxy substituents of the natural product isotrilobine (**2**) was realised to obtain two phenol groups that were substituted with the corresponding alkylamines to afford compounds **125-129** (**Figure 3A**)

Out of 5 new molecules tested, compound **125** showed significantly lower human plasma protein binding (90% in human plasma and 95% in mice). Importantly, this compound showed a similar antimalarial activity (123nM) as compound **84** (130nM), a good microsomal stability (half-life >60min) and an acceptable cytotoxicity. Moreover, the citrate salt of this compound is soluble in water, since no aggregation was found by Dynamic Light Scattering at a concentration of 15mM (**Figure 3B**).

### *In vivo* studies of 125

Thus compound **125** was chosen for *in vivo* studies. First, pharmacokinetic properties were assessed (**Figure 3C**): **125** has a long half-life (superior to 24h) with very slow elimination after a single dose administered either *i*.*v*., *p*.*o*. or *i*.*p*.. Plasmatic concentration reaches up to 600 ng/mL, 30min after *i*.*p*. injection. Antimalarial *in vivo* activity was then conducted using the Peter’s 4-days suppressive test in C57BL/6 mice infected *i*.*v*. with *P. berghei* ANKA (**Figure 3D**).

While the lowest dosage at 10mg/kg *i*.*p*. increased the survival by 3.5 days compared to the control, it did not reduce parasitemia. Dosing at 20mg/kg *i*.*p*. increased survival by 5 days and significantly lowered parasitemia compared to the control group (two-way ANOVA with Dunett’s correction, adjusted *p* value<0.0001). We didn’t increase further the dosage due to the potential toxicity of this compound (cytotoxicity around 2μM and slow elimination).

### Activity of 125 in the transmission and liver stages

First, we tested compound **125** in early (stage IIb-III) and late (stage IV-V) gametocytes stages (**Figure 4**). This compound has an IC_50_ around 1μM in both stages in *P. falciparum* NF54 gametocytes and was slightly more active in the Cambodian strain 3601E1 resistant to artemisinin (Kelch13 R539T), with an IC_50_ of 700nM (**Figure 4B**). Interestingly, the IC_50s_ were identical in both early and late gametocyte developmental stages, while antimalarials are usually less efficient against stage V, that is in a temporary quiescent state. For example, puromycin, used here as reference (**Figure 4B**), is less active against late stages, with curves that do not reach complete killing. We then assessed the efficacy of parasite development in the *Anopheles* mosquito stages by adding **125** to the infectious blood meal. The effect on parasite transmission was evaluated by counting the number of midgut oocysts, 10 days after the blood feeding (**Figure 4C**). Compound **125** did not increase mosquito mortality compared to DMSO (data not shown) and it decreased significantly the oocyst number at 10μM. Lastly, we tested compound **125** in primary human hepatocytes infected with *P. falciparum* sporozoites. Sporozoites invasion was performed for 3h, then cells were washed to remove remaining sporozoites and treated with **125** for 72h. Parasite survival was assessed by counting the number of trophozoites using immunofluorescence staining with anti-HSP70 antibody. We observed inhibition of parasite development in hepatocytes, indicating that **125** targets proteins expressed during multiple life cycle stages of *P. falciparum*.

**Figure 4.**
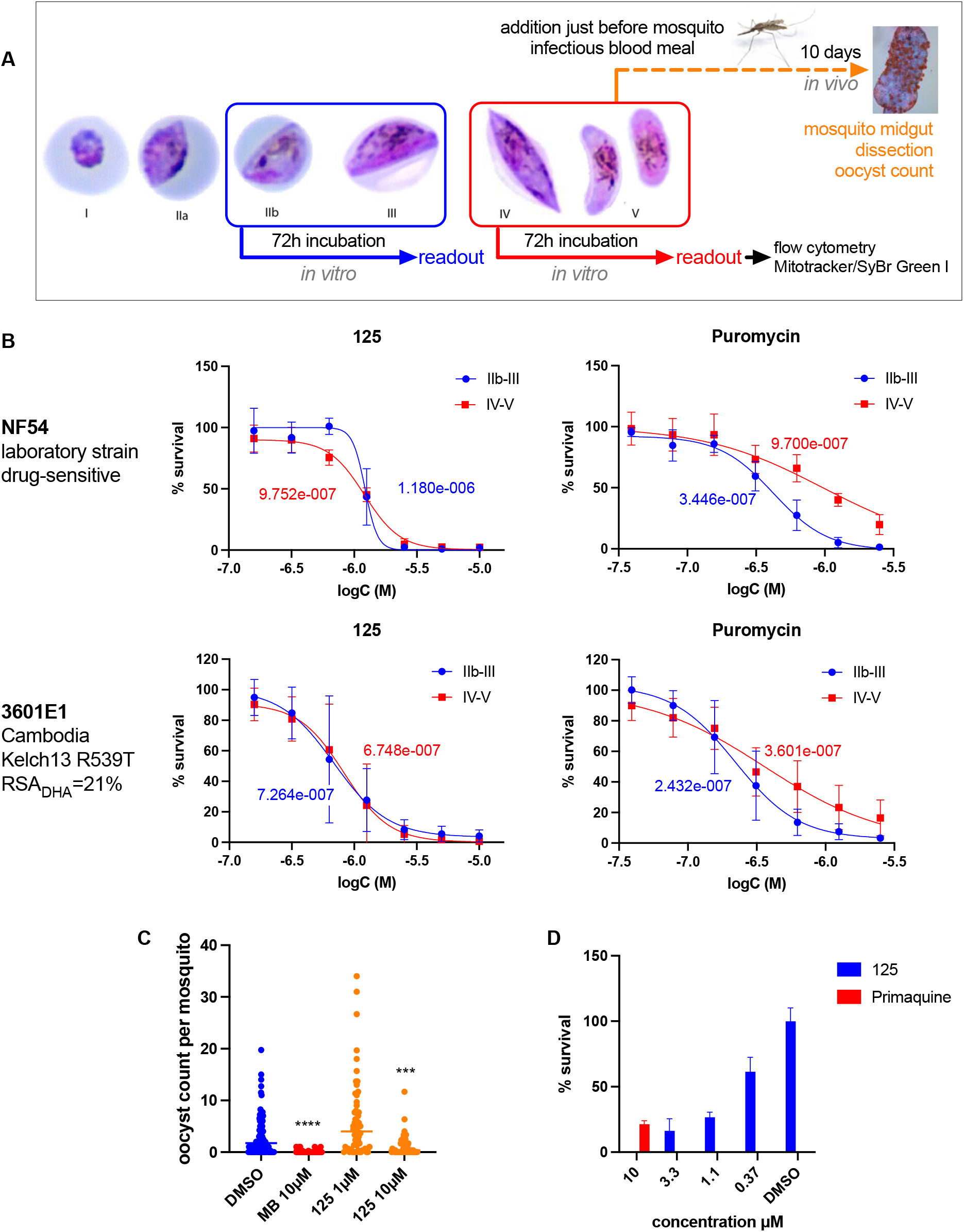
Activity of compound 125 in transmission and liver stages. **A**. Scheme showing the methodology used to assess anti-transmission activity. Early (IIb-III, blue) and late (IV-V, red) stage gametocytes were treated *in vitro* for 72h before survival was assessed using a flow-cytometry method (SyBR-Green I and Mitotracker deep red FM dyes). For *in vivo* mosquito studies (represented in orange), treatment was added to the infectious blood meal, containing mature stage V gametocytes, just before mosquito feeding. After 10 days of development, the number of oocysts was counted in each mosquito midgut. **B**. Gametocytocidal activity of **125** against early (IIb-III) and late (IV-V) stage gametocytes in NF54 drug-susceptible strain and 3601E1, a cloned Cambodian isolate bearing a R539T Kelch13 mutation that confers 21% survival after exposure to 700nM dihydroartemisinin (DHA) in the Ring-stage Survival Assay. Puromycin was used as positive control. N=3 independent experiments run in triplicates. **C**. Impact of compound **125** on parasite development in the mosquito. Each dot represents the number of oocysts counted in each mosquito midgut. N=4 independent experiments. Statistical analysis was done using Mann-Whitney test. \*\*\*\**p* value <0.0001; \*\*\**p* value=0.0002. **D**. Activity of **125** in *P. falciparum* liver stages. Compound was added after sporozoites invasion of human primary hepatocytes. Media containing increasing concentration of the compound was changed daily for 72 hours. Primaquine at 10μM was used as a positive control. Parasite survival was assessed using an immunostaining of parasites with *Pf*HSP70 antibody. N=1 experiment run in triplicate.

### Investigation of *P. falciparum* cellular targets

To decipher the mode of action of this bisbenzylisoquinoline alkaloid, we synthetized a chemical probe **132** derived from the parent natural products trilobine (compound **1**) and harboring a photo-activable moiety and an alkyne group (**Figure 5A**). It was used to photo-crosslink directly in cells proteins bound to the probe or in its vicinity and, by 1,3-dipolar cycloaddition, to functionalize the probe with a fluorescent reporter (TAMRA) and biotin for detection and pull-down studies followed by mass spectrometry analysis^9^. The probe **132** has a similar IC_50_ as trilobine (790nM versus 410nM) and was then used to treat parasite either alone or in competition with increasing concentrations of the parent, trilobine. Harvested parasites were saponin lysed and UV-irradiated at 365nm to crosslink the compound with the nearby target. Cells were then lysed and click reaction was initiated in parasite extracts. Purified functionalized proteins were then enriched using neutravidin beads and sent for analysis by mass spectrometry (**Figure 5A**). In gel fluorescence of the different conditions is showed in **Figure 5B**. Target deconvolution was done by plotting the competition curves *i*.*e*. protein abundance when the probe **132** was used alone (10μM) or in competition with 25μM, 50μM or 100μM of parent compound **1**. Only candidates producing curves with consistency between triplicates were selected. Among these, proteins that were not enriched compared to DMSO were discarded, as well as proteins with shared peptides (multigenic families). Clustering of the selected proteins was done using STRING software^10^ and clustered 2 different pathways: polyadenylate-binding protein 1 complex, involved in protein translation, and proliferating cell nuclear antigen involved in DNA replication (**Figure 5C and D**). The results indicate that multiple proteins interact with this family of compounds and, importantly, none of those candidates has been previously associated to drug targeting by antimalarials or drug-resistance mechanism.

**Figure 5.**
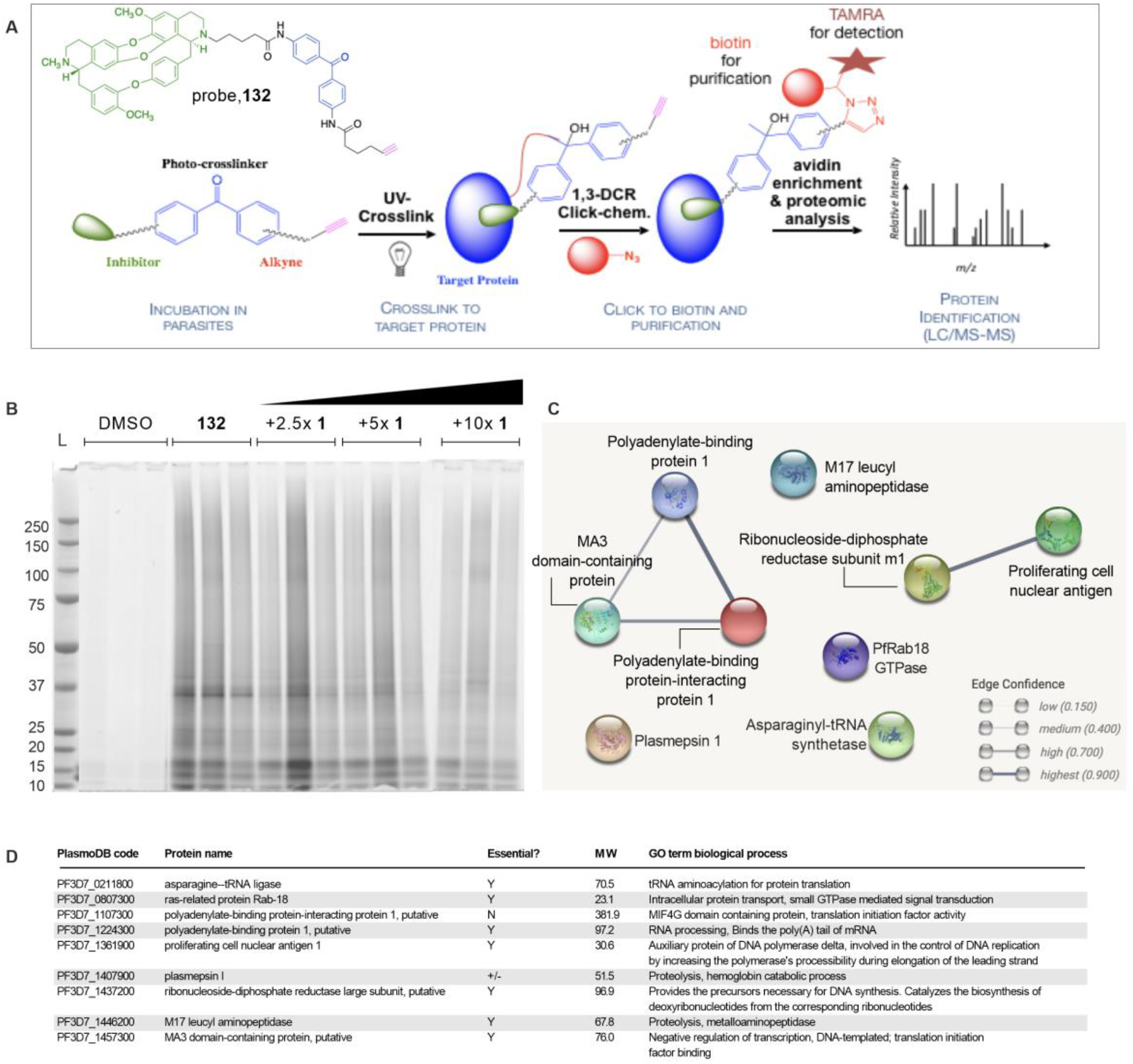
Chemical pull-down to decipher the protein targets. **A**. Chemical pull-down strategy. Chemical probe **132**, containing a photo-activable crosslinker moiety (in blue) and a reactive alkyne (in purple) to initiate the bioorthogonal 1,3-cycloaddition reaction, is incubated with parasites and binds to the protein partner(s). After photo-crosslinking, click-labelling with biotin and a fluorescent reporter (TAMRA) and avidin-biotin affinity purification, enriched protein partners are analyzed by mass spectrometry. **B**. In-gel fluorescence of the pulled-down proteins stained with TAMRA. The probe **132** was incubated with parasites at a concentration of 10μM and competition with trilobine (**1**) was done at 25μM (2,5x), 50μM (5x) and 100μM (10x), in triplicate. DMSO was used as a negative control. **C**. Clustering of enriched protein selected for their (i) decreasing quantity in competition with trilobine, (ii) enrichment compared to DMSO, and (iii) consistency between triplicates. Clustering was realized using STRING^10^ database **E**. Table reporting the essentiality^38^, molecular weight (MW) and GO term biological process of the chemical pull-down enriched proteins.

## Discussion

We explored the activity of bisbenzylisoquinoline alkaloids isolated from the plant *Cocculus hirsutus* on *P. falciparum*. The antimalarial activity of trilobine and its natural alkaloids analogues was significantly improved by synthesizing a library of hemisynthetic compounds. Importantly, none of the multi-drug resistant clinical isolates from Cambodia showed cross-resistance to the lead compounds, indicating that this chemical scaffold targets parasite pathways distinct from commonly used antimalarial drugs. Furthermore, we show that our lead compounds are fast-acting compounds, comparable to chloroquine and artemisinin. Another important activity is that the lead molecule kills early-ring stage, which is able to resist to high concentration of artemisinin by entering into dormancy^11^. The lead compound **84** can restore antimalarial activity in artemisinin-resistant isolates, when combined with DHA (**Figure 2C**), which highlights the potential of this chemical series to be used in combination with artemisinin. Natural bisbenzylisoquinoline alkaloids extracted from diverse plants have been already described for their antimalarial properties^12–18^, but no hemisynthetic studies have been reported to improve the pharmacological properties of these natural products. Previous re-sensitization has been described for bisbenzylisoquinolines in chloroquine-resistant *P. falciparum* strains, as well as in tumor cells resistant to vinblastin^16,19^.

As compound **84** was found to be highly bound to plasma protein (>99%), precluding its use for *in vivo* studies, new derivatives were synthesized resulting in compound **125** with a decreased plasma protein binding. Compound **125** showed sustained antimalarial activity (123nM), water solubility, high microsomal stability, and reasonable cytotoxicity (**Figure 3B**). Pharmacokinetic studies revealed rapid absorption and very slow elimination with an elimination half-life far superior to 24h (**Figure 3C**). *In vivo* antimalarial activity in mice infected with *P. berghei* shows that four doses at 20mg/kg of **125** delays the onset of parasitemia and increase mice survival by 5 days (**Figure 3D**). However, complete cure was not achieved, and dose escalation was avoided due to potential toxicity in mice at higher doses.

Compound **125** is also targeting transmission stages (*in vitro* and *in vivo*) and liver stages of *P. falciparum*, though at higher concentration than in the blood stage (**Figure 4**). This indicates that the protein target(s) are expressed in different stages of *Plasmodium*’s life cycle. To decipher the targeted proteins, we synthetized a chemical probe from the parent compound trilobine and used UV-affinity capture and chemical pull-down followed by mass spectrometry analysis. Target deconvolution of mass spectrometry hits revealed different protein interactions with the probe. The predicted targets have in common to have house-keeping functions in diverse vital pathways of *P. falciparum* such as the polyadenylate binding protein 1 (PABP1, Pf3D7_1224300) and the proliferating cell nuclear antigen 1 (PCNA1, Pf3D7_1361900) (for details, see **Figure 5D**). PABP1, its protein partner polyadenylate binding protein interacting protein 1 (PABP-IP1, Pf3D7_1107300) and MA3 domain-containing protein (MDCP, Pf3D7_1457300) are involved in protein translation^20,21^. They bind to the polyA tail of mRNA and are part of a complex with Eukaryotic translation Initiation Factor 4F (EIF4F) to mediate the recruitment of ribosomes to mRNA^22^. The PCNA1 protein is involved in DNA replication, by increasing DNA-polymerase delta processivity^23^, and in DNA-damage response^24^. PCNA1 is clustering with the ribonucleoside-diphosphate reductase subunit m1 (RNRm1, Pf3D7_1437200). This protein has an important role in DNA synthesis since it catalyzes the rate-limiting step of *de novo* synthesis of the deoxyribonucleotides by reducing the corresponding ribonucleotides^25^. To our knowledge, the identified pathways have never been targeted by antimalarials or have been involved in a malaria parasite drug-resistance mechanism. Interestingly, these pathways are vital for the other life cycle stages targeted by these compounds (transmission, liver stages). We show here for the first time that the bisbenzylisoquinoline trilobine could affect specific proteins involved in DNA replication and protein translation. This work lays out strategies for further studies to validate the protein target(s) by complementary methods such as thermal shift assays and the use of recombinant proteins. Defining the target will help the future preclinical drug-development process, by facilitating the improvement of selectivity compared to human homologues protein(s).

In conclusion, we identified a novel chemical scaffold derived from a natural product with activities against all life cycle stages that interacts with diverse essential parasite pathways.

## Methods

### Chemical library synthesis

Purification of trilobine and its natural products and hemi-synthesis of the trilobine derivatives are described in^5,26^.

### Compound solubility/aggregation

Compounds were mixed with citrate (1:1) to obtain the salt and then diluted in water at a final concentration of 15mM. 20μL of each sample was then screened for aggregates by Dynamic Light Scattering, on a Dniproplate reader III (Wyatt technology).

### Microsomal stability and plasma binding

Rat and human microsomal stability and plasma binding studies were outsourced at Fidelta Ltd., Zagreb, Croatia.

### Cytotoxicity in the HepG2 and MV411 cell lines

Human HepG2 (hepatocellular carcinoma, DSMZ) and MV411 (lymphoblasts, acute monocytic leukemia, DSMZ) cells were grown as advised by the provider. Cytotoxicity was evaluated after 72h of drug treatment in 96-well plates by ATPlite (PerkinElmer) according to the manufacturer’s protocol. Experiments were run in technical triplicates and at least in two biological replicates. GraphPad Prism 8 was used to interpolate IC_50_. DMSO was used as control.

### *P. falciparum* Continuous Culture

*P. falciparum* parasites were cultured using a standard protocol^27^. The laboratory strains used were NF54 and 3D7; *P. falciparum* clinical isolates are described in the relevant section.

### Inhibition Activity in the Asexual Stages

IC_50_ values were obtained as previously described^28^. Briefly, a range of 7-point and 2-step serial dilutions starting at 2μM were used to assess the activity of the compounds. GraphPad Prism 8 was used to interpolate IC_50_ from three independent experiments run in triplicate. DHA and DMSO were used as positive and negative controls, respectively.

### Stage-Specificity during the Asexual Cell Cycle

Stage-specificity was assessed using previously described methods^28^. Briefly, asexual NF54 parasites were tightly synchronized (0–3hpi) and incubated for 6h with compound **84** at 3× and 10× IC_50_ value, at 0, 6, 12, 18, 24, 30, 36, or 42hpi. Following each treatment, cells were pelleted, washed with 10mL of RPMI, and put back into culture in a new plate. Parasitemia was assessed at 72h post-synchronization using Giemsa-stained thin blood smears. The percentage of survival was compared to DMSO-treated parasites. Data were obtained from three independent experiments (one well per condition).

### *In vitro* Parasite Reduction Rate (PRR)

The *in vitro* parasite reduction rate was adapted from Sanz et al^7^. 10^6^ infected RBC containing mostly rings were cultivated separately with 10× the IC_50_ of **84** or reference drugs (DHA, Chloroquine or Pyrimethamine) for 6h, 24h, 48h, 72h and 96h (0.5% starting parasitemia, 2% hematocrit of *P. falciparum* 3D7). After each incubation time, cells were washed and serially diluted (⅓) with fresh RBC. Growth was assessed after 3 weeks, using the SYBR-green I assay. The number of wells showing parasite growth is correlated to the number of viable parasites at the different time points (as defined by Sanz *et al*.) and allows defining the growth parameters reported in **Figure 2B**. The experiment was performed once in duplicate.

### Antiplasmodial activity against *P. falciparum* field isolates

*P. falciparum* isolates used in this study were collected in the framework of the therapeutic efficacy surveillance program in Cambodia between 2017 and 2019. PSA, RSA, AQSA and genotyping data have been performed in the Pasteur Institute in Cambodia. Parasites were chosen based on the pattern of multi-drug resistance they carry. IC_50_ values were obtained as previously described^28^ with slight modifications. Ring-synchronized cultures (2% hematocrit, 1% starting parasitemia) were incubated for 96h before growth measurement using the SYBR-green I assay. A range of 11-point and 2-step serial dilutions, starting at 1μM were used to assess the activity of **84**. Mefloquine (MQ) and monodesethylamodiaquine (dAQ) were used as positive controls. Experiments were performed in triplicate, with one biological replicate per isolate. Statistical analysis was performed using multiple Mann-Whitney tests in comparison with the *drug-sensitive Cambodia* condition.

#### Ring-stage Survival Assay (RSA)

The RSA was determined as previously described^8^.8/30/2022 9:52:00 PM Briefly, 0-3h synchronized ring-stage parasites were exposed to a 6h treatment with either DMSO, 700nM DHA, 3xIC_50_ of **84** or the combination of both. The IC_50_ used was the one obtained against the NF54 strain. After 6h, drugs were washed out and parasites were put back into culture. Parasitemia was assessed at 72h post-synchronization using giemsa-stained thin blood smears. Percentage of survival was compared to DMSO-treated parasites and data were obtained from one independent experiment, in the different isolates. Statistical analysis was performed using multiple Mann-Whitney tests in comparison with the *drug-sensitive Cambodia* condition.

### Pharmacokinetic studies

PK studies were outsourced at Fidelta Ltd., Zagreb, Croatia, according to standard procedures.

### *In vivo* antimalarial activity

All animal experiments performed in the manuscript were conducted in compliance with French animal welfare laws and Institut Pasteur Animal Care and Use Committee guidelines. *In vivo* activity was determined as previously described^29^ following the Peters 4-day suppressive test^30^ with slight modifications. 6-weeks old C57BL/6 female mice (Janvier Labs) were infected intravenously (*i*.*v*.) with 10^5^ iRBC of *P. berghei* ANKA GFP-expressing parasites^31^. 2h post-infection, mice were treated intraperitoneally (*i*.*p*.) for 4 days with **125** following a daily regimen of 20mg/kg or 10mg/kg, or 25mg/kg of chloroquine, or the equivalent vehicle control. Parasitemia was quantified from blood samples collected every day by flow cytometry of 200,000 RBCs and confirmed by Giemsa-stained thin blood smears. The experiment was performed once (n=6 mice per group).

### Activity against synchronous stage IIb-III and stage IV-V gametocytes

For this experiment, we used two different strains that retain their capacity to produce gametocytes: the drug-susceptible laboratory strain NF54 and an artemisinin-resistant (Kelch13 R539T) cloned Cambodian isolate 3601E1. Synchronous gametocytes were induced following previously described methods^32^ with slight modifications. Briefly, trophozoite purification steps were performed using gelatin flotation (Plasmion®). Media containing 0.05% (w/v) Albumax I was used until induction (day 0, corresponding to gametocyte-committed ring stage parasites), followed by a media containing 5% (v/v) human serum and 0.025% (w/v) Albumax I. Asexual parasite populations was removed by *N*-acetyl Glucosamine treatment for 5 days. Parasites were treated with **125** and the positive control puromycin to evaluate efficacy against early (day 4, stage IIb-III) and late-stage gametocytes (day 9, stage IV-V). Parasites, at approximately 1% starting gametocytemia, were seeded in 96-well plates at 2% final hematocrit and treated for 72h with a 7-point and 2-step serial dilutions of compounds starting at 10μM for **125** or 2.5μM for puromycin. Readout of parasite survival was done by flow cytometry (Guava 6HT®, Luminex corp) using a Mitotracker® Deep Red^FM^ (molecular probes, 50μg, reference M22426) and SYBR™ Green I (10,000x concentrate in DMSO, Lonza) staining. SYBR Green I was used at a final dilution of 0.5x and Mitotracker at a final concentration of 100nM, in prewarmed PBS (37°C). Data were acquired on 10,000 red blood cells, in triplicate, in 3 independent experiments.

### Standard membrane feeding assay

The culture of gametocytes is performed in a semi-automated system developed in Nijmegen, Netherlands^33^ at an initial parasitemia of 0.5%. After 4-6 days, the parasitemia reaches approximately 5-10 % and gametocytogenesis is induced. The cultures are checked twice a week to evaluate the presence of different stages of gametocyte development. Infectious stage V gametocytes are observed after approximately two weeks of culture. To assess the maturity of the male gametocytes, an aliquot of the culture is taken and the number of exflagellation events are counted under optical microscope, using a 400x magnification, by diluting infected red blood cells in human serum. Cultures containing mature gametocytes are diluted to 1:6 in a mix of 50% fresh red blood cells and 50% human serum. Compound **125** (1 or 10μM final concentration), methylene blue (10μM) or DMSO are added to this mix and fed to 50 *Anopheles stephensi* females per condition, using a membrane feeder. The mosquitoes are maintained at 26°C and 70% humidity with 10% sucrose as food source. Ten days post-infection, the mosquito midguts are dissected and stained with 0.25% fluorescein and the number of oocysts per midgut are counted. The results have been obtained in 4 independent experiments. Statistical analysis was done using Mann-Whitney test.

### Liver-stage activity

Micropatterned human primary hepatocytes surrounded by supportive stromal cells^34^ were seeded on glass-bottom 96-well plates and infected the following day with 70,000 sporozoites per well of *P. falciparum* NF54. Cells were centrifuged at 700g for 5min to settle down the parasites. Sporozoites were allowed to invade for 3h before washing the cells, then fresh media containing drugs was added and changed daily. Cells were incubated for 72h at 37°C and 5% CO_2_ before fixation and permeabilization with ice-cold methanol and staining with dapi and PfHSP70 antibody (StressMarq BiosciencesINC., catalog number SPC-186). Readout was done by counting PfHSP70-stained exoerythrocytic forms in fluorescence microscopy. **C-125** was tested at 4 different concentrations, ranging from 10 to 0.37μM, using a 3-fold dilution. Primaquine diphosphate at 10μM was used as a control. Experiment was done once in triplicate.

### UV-affinity capture and protein pull-down

Asynchronous *P. falciparum* NF54 cultures (1.10^8^ parasites per condition) were treated for 1 hour with 10μM of probe or DMSO. After 30min competitor was added in increasing concentration (DMSO, 25μM, 50μM or 100μM of trilobine, **1**) for 30min before parasites were harvested, saponin-lysed in ice (0.15% saponin in PBS) and washed thoroughly in ice-cold PBS to remove hemoglobin contaminants. Parasite pellets were resuspended in PBS and irradiated for 1h at 365nm, on ice, to crosslink the probe to the protein target. Parasites were centrifuged and pellets were lysed in a buffer containing 1% SDS, 1mM dithiothreitol (DTT) and protease inhibitors (Roche) in PBS. Protein concentration was determined and extracts were diluted to a concentration of 1mg/ml. Click-labelling was done following described protocol^35^. Briefly, extracts were treated with 6μL/100μL of extract of a premix solution containing: 60μM of TAMRA-biotin-azide (Jena Bioscience, catalog number CLK-1048-5), 1mM CuSO_4_ in water (freshly prepared), 1mM tris(2-carboethyl)phosphine (TCEP) and 100μM tris[(1-benzyl-1*H*-1,2,3-triazol-4-yl)methyl]amine (TBTA). Tubes were shaken at room temperature for 1h after which proteins were precipitated with sequential addition and vortexing of methanol (200μL), chloroform (50μL) and ddH_2_O (100μL). Tubes were then centrifuged for 2min at 14,000g and protein layer was washed twice with 200μL of methanol. Methanol was discarded and remaining solvent was let to evaporate at room temperature. Proteins were resuspended in SDS buffer (1% SDS, 10mM DTT). Samples were then enriched on avidin-coupled agarose beads (Pierce™NeutrAvidin™ Agarose, ThermoFisher Scientific; 50μL, pre-washed three times in 0.2 % SDS in PBS) by incubation with gentle shaking for 2h at room temperature to bind and enrich biotin-labeled proteins thanks to the probe. The supernatant was then removed, and the beads were washed on a column with 1% SDS in PBS (3 × 500μL), 4M urea in 50mM ammonium bicarbonate (AMBIC) (2 × 300μL) and 50mM AMBIC (5 × 300μL). This experiment was done in three independent replicates.

### Protein digestion

Reduction of disulfide bonds of enrich biotin-labeled proteins in 50mM AMBIC was achieved by addition of DTT to final concentration of 5mM and incubation for 30min at 56°C with shaking. Then, IAA was added at final concentration of 20mM, and samples were incubated in dark for 30min at 37°C. Samples were digested for 4h using LysC (Promega, MA, USA). Subsequently, 1μg of trypsin (Promega, MA, USA) was added, and samples were digested overnight at 37°C. Digestion was quenched by addition formic acid (FA). The digest was subsequently desalted using Sep-Pak C18 cartridges (Waters) and lyophilized in vacuum for further analysis.

### LC-MS/MS analysis

All the samples were analyzed on a Q Exactive™ Plus Hybrid Quadrupole-Orbitrap™ Mass Spectrometer coupled with an Easy nLC 1200 ultra-high pressure liquid chromatography system (Thermo Fisher Scientific) with solvent A of 0.1% formic acid and solvent B of 0.1% formic acid/80% acetonitrile. One microgram of each sample was injected on a 75-μm ID PicoFrit column packed *in-house* to approximately 30-cm length with ReproSil-Pur C18-AQ 1.9-μm beads (Dr. Maisch). Column equilibration and peptide loading were done at 900 bars in buffer A (0.1% FA). Samples were separated at a 250nL/min flow rate with a gradient of 2 to 7% buffer B in 5min, 23% buffer B in 70min, 23 to 45% buffer B in 30min, 45 to 95% buffer B in 5min, followed by a hold at 95% for 7min and back to 2% for 15min. Column temperature was set to 60°C. Mass spectrometer was operated in data-dependent acquisition mode. The MS instrument parameters were set as follows: MS1, r=70,000; MS2, r=17,500; MS1 AGC target of 3e6; MS2 for the 10 most abundant ions using an AGC target of 1e6 and maximum injection time of 60ms; and a 45s dynamic exclusion. The isolation window was set to 1.6m/z and normalized collision energy fixed to 28 for HCD fragmentation. Unassigned precursor ion charge states as well as 1, 7, 8 and >8 charged states were rejected and peptide match was disable.

### Data analysis

Acquired raw data were analyzed using MaxQuant 1.5.3.8 version^36^ using the Andromeda search engine^37^ against *Plasmodium falciparum*, strain NF54 proteome database (5,548 entries, download from PlasmoDB) and human reference proteome (89260 entries, download Jan2018), concatenated with usual known mass spectrometry contaminants and reversed sequences of all entries. All searches were performed with oxidation of methionine and protein N-terminal acetylation as variable modifications and cysteine carbamidomethylation as fixed modification. Trypsin was selected as protease allowing for up to two missed cleavages. The minimum peptide length was set to 7 amino acids. The false discovery rate (FDR) for peptide and protein identification was set to 0.01. The main search peptide tolerance was set to 4.5ppm and to 20ppm for the MS/MS match tolerance. One unique peptide to the protein group was required for the protein identification. The match between run was selected. A false discovery rate cut-off of 1% was applied at the peptide and protein levels. All mass spectrometry proteomics data have been deposited at ProteomeXchange Consortium via the PRIDE partner repository with the dataset identifier PXD036288. The identification and quantification data were then mined for functional analysis to identify targets using STRING^10^.

## Acknowledgements

We thank Bertrand Raynal of the Plateforme de Biophysique Moléculaire, in C2RT Pasteur Institute, for technical help. We thank the Center for Production and Infection of *Anopheles* (CEPIA) in Institut Pasteur for the mosquito breeding and handling.

We thank Bruno Vitorge and Remy Lemeur from the Institut Pasteur Biological NMR Technological Platform for assisting with NMR experiments, Frédéric Bonhomme of the CNRS-Institut Pasteur UMR3523 Organic Chemistry Unit for performing HRMS analysis.

This work was supported by Institut Pasteur-Institut Carnot (S-CR18089-02B15 DARRI CONSO INNOV 46-19; S-PI15006-10A INNOV 05-2019 ARIMONDO IARP 2019-PC), Pasteur Transversal Research Program (PTR 233-2019 HALBY), Pasteur Swiss Foundation grant, Agence Nationale de la Recherche (ANR EpiKillMal), and Pasteur-Roux-Cantarini Fellowship.

